# Diversity of vector-dispersed microbes peaks at a landscape-defined intermediate rate of dispersal

**DOI:** 10.1101/2025.11.17.688957

**Authors:** Yuta Nagano, Kae Natsume, Tadashi Fukami, Kaoru Tsuji

## Abstract

Dispersal rate has long been considered a primary determinant of species diversity in ecological communities. However, this knowledge is mostly built on studies of organisms that disperse actively by themselves or passively via physical forces. For organisms that disperse passively via other organisms, the landscape matrix that affects their vectors should indirectly shape the diversity of the vectored, but this relationship remains poorly understood. We investigated landscape–dispersal–diversity relationships in nectar-inhabiting bacteria that disperse via flower-visiting insects. Field observation and experiments revealed that bacterial diversity and abundance peaked at an intermediate frequency of insect visits, which was in turn determined by the surrounding landscape characteristics observed at the 200-m radius scale. Based on this finding, we discuss the possibility that species diversity tends to be maximized at an intermediate dispersal rate, especially when habitat patches of vectored organisms constitute consumable resources for their vectors.

## Introduction

Dispersal rate, or the rate at which individuals disperse from regional external sources to local habitat patches, has long been considered a primary factor shaping local biodiversity (MacArthur & Wilson 1967). Theory and empirical evidence suggest that diversity is highest under an intermediate rate of dispersal. As dispersal rate increases, the probability that not only new species but also dominant competitors colonize in local habitat patches increases, resulting in reduced diversity due to competitive exclusion under very frequent dispersal (Cadotte 2006; Mouquet & Loreau 2003). However, whether diversity shows a hump-shaped pattern with dispersal rate as predicted by this intermediate dispersal hypothesis (Cadotte 2006; Mouquet & Loreau 2003) or increases monotonically with dispersal rate (Loke & Chisholm 2023; Loreau & Mouquet 1999) can depend on the spatial scale of observation relative to that of local habitat patches (Lu et al. 2019). Furthermore, increasing attention has been paid to better understanding determinants of dispersal rate (Baguette et al. 2013).

Besides obvious differences among species that vary in their dispersal capacity being a factor, another consideration is the landscape matrix. Even the same type of organisms can vary greatly in dispersal rate, depending on, for example, how permeable the landscape surrounding the focal habitat patch is to the dispersing organisms (Mony et al. 2020, 2022).

These investigations into the relationship between dispersal, diversity, and landscape characteristics have often assumed that dispersal happens either actively as organisms move or passively with physical forces such as wind, rain, and gravity (Loke & Chisholm 2023; Vanschoenwinkel et al. 2013). For these organisms, the connectivity (e.g., distance) between the regional source pool and the local habitat patches largely determines dispersal rate (McArthur & Wilson 1967). However, a wide range of plants, animals, fungi, and bacteria disperse by hitchhiking on an organism of another species that moves a longer distance than they do themselves, the mode of dispersal known as phoresy (Bartolow & Agosta 2021).

Examples include ungulate-dispersed grasses (Plue & Cousins 2018), bird-dispersed fishes (Lova-Kiss et al. 2020), rodent-dispersed fungi (Borgmann-Winter et al. 2023), and fly-dispersed bacteria (Wohlfahrt et al. 2020), to name just a few (Bartlow & Agosta 2021). For these species that are carried biotically by dispersal vectors, the landscape matrix affecting the movement of the vectors, rather than the vectored, influences dispersal rate and, consequently, diversity. To understand landscape–dispersal–diversity relationships more comprehensively, the effects of vector-mediated dispersal should be studied in a landscape context. Yet, few empirical studies have addressed these effects.

In this paper, we report the results of a field study that examined how landscape matrix shapes the diversity of vector-dispersed organisms indirectly by affecting their vectors. Specifically, we focus on bacteria that inhabit floral nectar, which has been increasingly used as a natural microcosm (sensu Srivastava et al, 2004) to address fundamental questions in population and community ecology over the past decade (Chappell & Fukami 2018). These bacteria often rely on flower-visiting insects to disperse among flowers (Cusumano et al. 2023; Herrera et al. 2008; Vannette & Fukami 2017). We first sought to identify the characteristics of the surrounding landscape that affected the frequency of insect visits to flowers. Building on this analysis, we then investigated the relationship between the rate of insect-mediated bacterial dispersal and the diversity of bacterial taxa through a complementary pair of field observation and experiment. Both observational and experimental data revealed hump-shaped responses of bacterial diversity to the rate of insect-mediated dispersal as determined by landscape matrix. Informed by these findings, we discuss the possibility that this intermediate dispersal pattern is commonly observed in systems where habitat patches of the vectored organisms constitute consumable resources for their dispersal vectors.

## Materials and methods

### Study site

This study was conducted in September 2020 in buckwheat (*Fagopyrum esculentum*) fields embedded in a rural landscape consisting of a small-scale mosaic of agricultural fields, woodland, grassland, riparian zones, and residential areas in the town of Iijima in Nagano Prefecture, located in central Japan (**Figure 1A-C**, 35°67’67“N, 137°91’96“E). In this region, buckwheat is cultivated with the same amounts of seeds sown (0.4 – 0.5 kg/a) and fertilizers applied (100kg/a) across hundreds of small fields (many are less than 0.5 ha), making the fields comparable in these respects. No agrochemicals (e.g., pesticides, herbicides, fungicides) are used during buckwheat cultivation.

**Figure 1.**
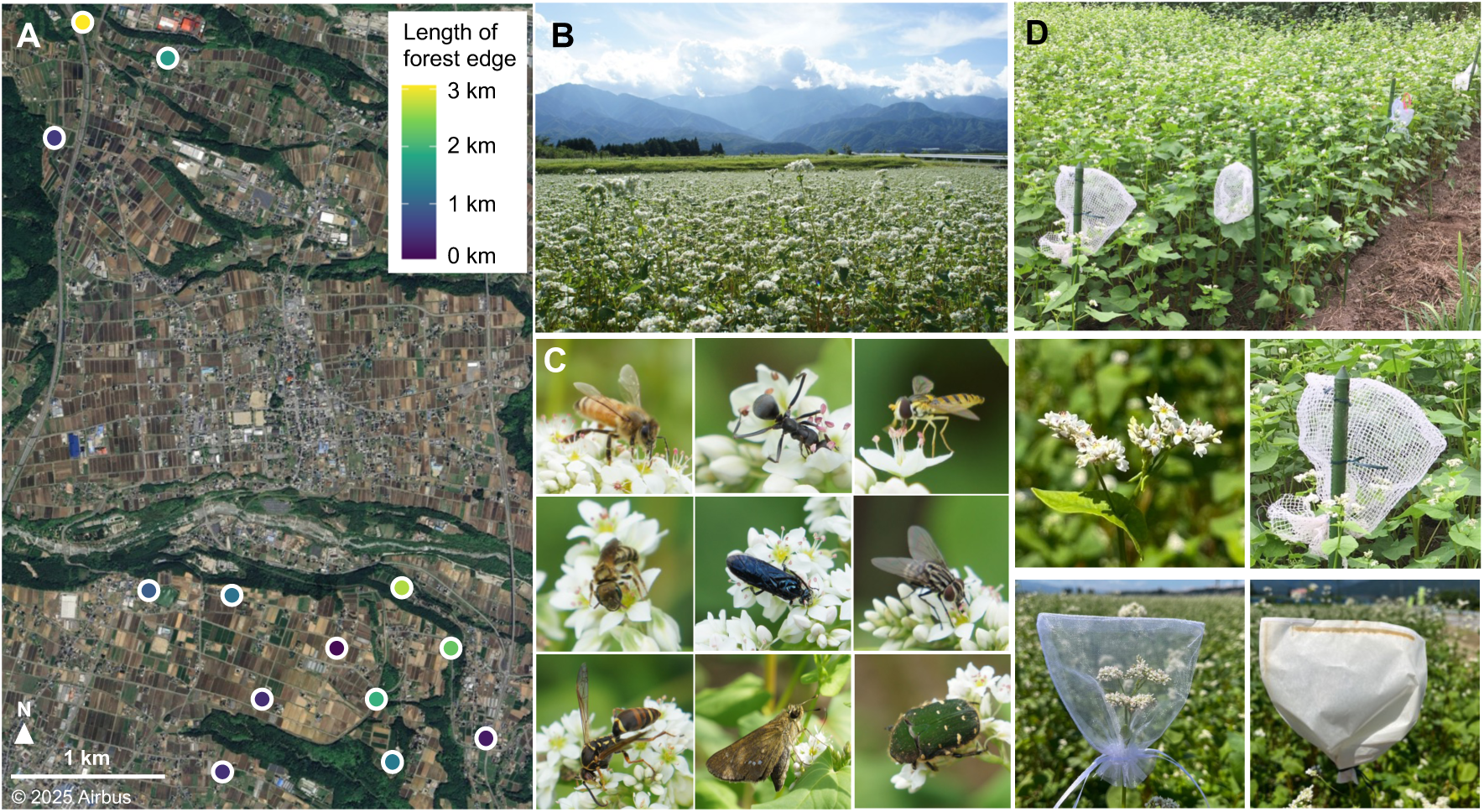
Study system and bagging methods. (A) Map showing the location of the 13 buckwheat fields used in this study. The color indicates the length of forest edges within 200-m buffers, with brighter colors denoting longer forest edge lengths. (B) One of the buckwheat fields. (C) Some of the diverse insects that visit buckwheat flowers. From top-left to bottom-right photos, *Apis mellifera* (Hymenoptera: Apidae), *Formica japonica* (Hymenoptera: Formicidae), *Sphaerophoria macrogaster* (Diptera: Syrphidae), *Halictus aerarius* (Hymenoptera: Halictidae), *Arge similis* (Hymenoptera: Argidae), Sarcophagidae sp. (Diptera), *Polistes snelleni* (Hymenoptera: Vespidae), *Parnara guttata* (Lepidoptera: Hesperiidae), and *Gametis jucunda* (Coleoptera: Scarabaeidae). (D) Buckwheat inflorescences bagged for insect exclusion. The middle photos show no bag control, allowing access to all arthropods, and rough-mesh bag treatment (mesh size = 4.5 mm), allowing access only to arthropods smaller than 4.5mm long, such as small hoverflies and ants. The bottom photos show fine-mesh bag treatment (mesh size = 0.4 mm), allowing access only to arthropods smaller than 0.4 mm long such as thrips and mites, and paper bag treatment, allowing access to no arthropods.

Bacterial communities that form within floral nectar can be delineated clearly, with flowers on the plant and more broadly within agricultural fields forming metacommunities in a spatially hierarchical fashion (Chappell & Fukami 2018; Rering et al. 2024). Furthermore, the dispersal vectors of nectar-inhabiting bacteria, i.e., flower-visiting insects, are diverse in buckwheat, with microbial dispersal rate varying with insect abundance and diversity (**Figure 1C**, Miyashita et al. 2023; Nagano et al. 2021). Previous work in this landscape (Miyashita et al. 2023) and elsewhere in Japan (Taki et al. 2010) has shown that the frequency of flower visits by pollinators is determined by the landscape matrix of buckwheat fields. Specifically, from previous work (Bartual et al. 2019), more insects are expected to visit buckwheat flowers when there are more forest edges in the landscape surrounding the field. These characteristics of bacteria in buckwheat nectar make them a suitable system for studying the effects of landscape matrix and vector-mediated dispersal on the diversity of the vectored organisms at multiple spatial scales.

Nectar frequently contains yeasts in addition to bacteria, and these two groups can affect each other strongly within nectar (Álvarez-Pérez et al. 2019). Our preliminary work indicated, however, that buckwheat nectar in our study landscape has few yeasts and is predominantly inhabited by bacteria (see **Supplement Text 1**). This skewed representation of bacteria over yeasts is commonly observed in plant species of short-lived flowers with short corolla (Vannette et al. 2021). Buckwheat flowers have short corollas, and most flowers last only a single day. For this reason, we focused on bacteria in this study.

We selected 13 buckwheat fields for field observation and experiment for this study in a way that maximized variation in landscape matrix across fields. All fields were located at least 400 m away from the nearest-neighbor field used in this study (**Figure 1A**). In addition, the fields were chosen to minimize spatial bias among the fields in terms of forest edge length, such that fields having similar forest edge length would not be spatially aggregated (**Figure 1A and 2B**). The focus of this study was to understand how the characteristics of the landscape matrix surrounding a buckwheat field affected the foraging behavior of individual insects as microbial vectors. As such, this study differs from those that aim to understand how landscape characteristics influence the entire population dynamics of insects in the landscape (e.g., Eeraerts et al. 2019; Ekroos et al. 2013; Nicholson et al. 2017), which often require observation at larger spatial scales than needed for the purpose of this study.

**Figure 2.**
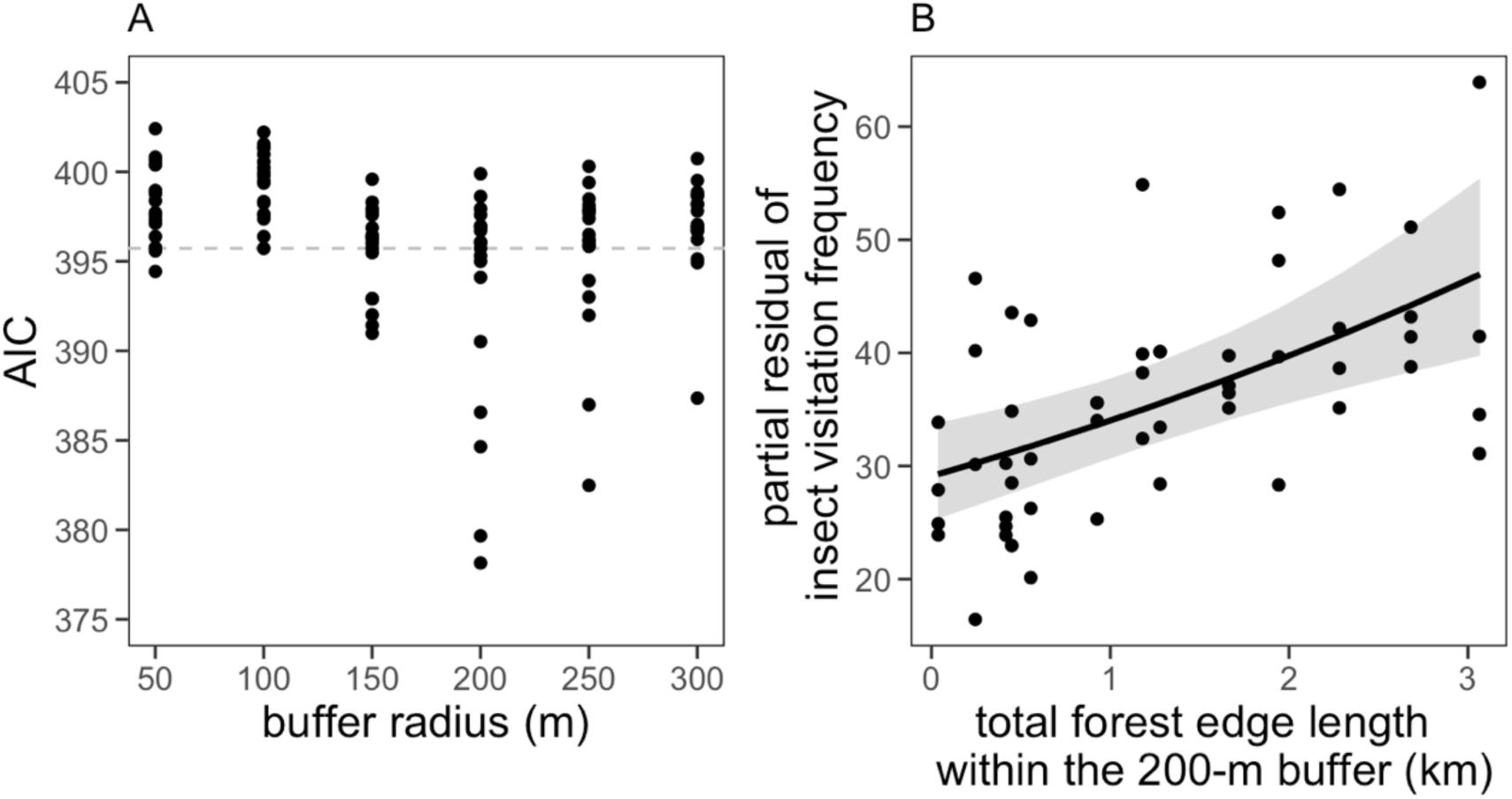
Relationship between landscape context and insect visitation frequency. (A) Results of AIC model selection, where each dot represents the AIC value of each model. Gray dashed line indicates the AIC value of the null model. (B) Relationship between landscape characteristics (as quantified by forest edge length) and the frequency of insect visitation to buckwheat flowers. Data points represent four observations at each of the 13 study fields, with the predicted mean and 95% confidence interval.

To quantify landscape matrix, we categorized human land use around fields into four types: forest edges, buckwheat fields, other agricultural lands (e.g., paddy fields and orchards), and residential areas. To calculate the area of each land-use type, buffers with a radius ranging from 50 to 300 m at 50 m intervals were generated from the edge of each field using GIS performed by R version 4.4.0 (R Core Team 2024) and sf package (Pebesma 2018). This range of buffer radius was based on previous reports indicating that a 100-300 m buffer radius best captures landscape characteristics for estimating the flower visitation frequency of insects, other than that of *Apis* and *Bombus* bees (Benjamin et al. 2013; Földesi et al. 2015; Steffan-Dewenter et al. 2002), both of which were rare in our study region in September (Miyashita et al. 2023, Nagano et al. 2021, 2025). The Shannon diversity index of land-use proportion was calculated as the landscape heterogeneity index. To characterize landscape matrix, forest edge length and landscape heterogeneity were used. We also recorded the management of field margin grasslands, i.e., whether mown or set-aside, because local field context, such as grassland management, can strongly influence flower-visiting insect densities in agricultural landscapes (Albrecht et al. 2020; Nagano et al. 2025). Vegetation height of wild plants in recently mowed field margins was less than 10 cm. Set-aside field margins that had not been mowed for at least one month had vegetation that was taller than 30 cm.

### Field observation

We used both observational and experimental approaches as complementary methods to investigate the relationship between the frequency of insect visits to flowers and the taxonomic diversity of bacteria in nectar. For the observational approach, we quantified the frequency of flower visits by insects at each of the 13 buckwheat fields (**Figure 1A**). For this purpose, we established four observational transects (1 × 30 m), with one transect placed along each of the four sides of the fields in most cases. In each observational transect, insects visiting buckwheat flowers were observed between 9:00 am and 11:50 am on four sunny days, including September 9, 14, 15, and 17 of 2020. This data collection involved walking along the field margin for three minutes per observation, following the method by Miyashita et al. (2023). Insect visitation frequency was recorded separately for ten insect groups: honeybees, wild bees, ants, wasps, sawflies, hoverflies, other flies, butterflies, flower chafers, and other beetles (**Figure 1C**, Nagano et al. 2025).

To quantify bacterial abundance and diversity in floral nectar, we established four sites at each of the 13 buckwheat fields (**Figure 1A**), with one site placed along each of the four sides of the field in most cases. At each site, we sampled nectar from two plants of different floral morphs, one pin and one thrum (Cawoy et al. 2006). From each plant, we collected nectar from four flowers on an inflorescence in the late afternoon between 4:00 pm and 5:00 pm on September 5, 8, 9, and 11-13 of 2020. Nectar was collected from four flowers on each of the inflorescences using a 0.5-µl microcapillary tube. The nectar from the four flowers was consolidated to make one sample. The total volume of each of these nectar samples was measured, and each sample immediately placed into a separate PCR tube containing 40 µl of distilled water. The diluted samples were kept cool in a Styrofoam box with ice packs until transported to the laboratory.

Within two hours of the sampling in the field, 5 µl of each diluted sample was then diluted further in the laboratory and plated onto tryptic soy agar (TSA; 1.5% tryptone, 0.5% soytone, 0.5% NaCl, 5% fructose, 1.5% agar, all (w/v)) plates with the antifungal cycloheximide to obtain CFU (colony-forming units) counts. We used the CFU counts as an estimate of bacterial abundance in nectar, which should be interpreted with caution as not all bacterial taxa may form colonies on any given media. However, many of the dominant bacteria taxa in floral nectar, such as species of *Acinetobacter*, *Rosenbergiella*, *Erwinia*, *Pantoea*, *Pseudomonas*, and *Sphingomonas*, are culturable on TSA (Quevedo-Caraballo et al. 2025; Vannette et al. 2021). The CFU counting was done after two days of incubation at room temperature (about 20-26 °C). The remainder of the original samples were stored at −80°C until used for DNA extraction and PCR for metabarcoding (**Supplementary Text 2**). The observed numbers of ASV_97_ for all nectar samples were in the range of saturation with respect to the read number (**Figure S1**).

### Field experiment

For the experimental approach to examine the relationship between insect visitation frequency and nectar bacterial diversity, we established four experimental blocks at each of the 13 buckwheat fields (**Figure 1A**). In most fields, we had one block along each of the four sides of the field. In each experimental block, we had two sets of the following three inflorescence treatments: an inflorescence covered with a fine-mesh bag (0.4 mm), an inflorescence covered with a rough-mesh bag (4.5 mm), and an inflorescence without a bag (control) (**Figure 1D**). We chose a pair of plants with different floral morphs for each treatment. The control inflorescences were the same as those used for field observation described above. In addition, in one of the four experimental blocks in each field, we had a fourth treatment, which was two inflorescences (one pin and one thrum) each covered with a paper fruit bag. The bagging was used to prevent insects of differing sizes from visiting the flowers on the bagged inflorescences. The paper bag, the fine-mesh bag, and the rough-mesh bag were intended to exclude all arthropods, allow only arthropods smaller than 0.4 mm long (e.g., thrips and mites), and allow only arthropods smaller than 4.5 mm long (e.g., small hoverflies and ants), respectively. The bagging was done in the morning between 8:00 am and 10:00 am (**Figure 1D**). Nectar collection, plating, DNA extraction and PCR for metabarcoding were done following the same methods as described above for field observation (**Supplemental Text 2**). We note that, as usual with any other systems, reduced diversity may sometimes represent reduction in population size below the detection threshold rather than true extinction.

The bagging treatment was used to experimentally create a gradient of insect visitation frequency, with no-bag treatment allowing for the highest frequency, followed by rough-bag, fine-bag, and paper-bag treatments in that order. The bagging treatment also caused changes in the types of insects visiting flowers, in addition to altering overall visitation frequency.

Although these effects can complicate interpretation of the experimental results, we found no significant difference in bacterial taxonomic composition among the four bagging treatments (**Figure S2**, PERMANOVA, p = 0.996), suggesting that no significant difference existed in the taxonomic identity of bacterial taxa vectored by insects of different sizes (see also **Figure S3**). Therefore, in our system, any difference in bacterial diversity we observed among the bagging treatments could be attributed mainly to insect visitation frequency rather than to insect species identity.

### Statistical analysis

All statistical analyses were conducted using R version 4.4.0 (R Core Team 2024). First, spatial autocorrelation for insect visit frequency among fields was checked based on Moran’s I statistics using xy coordinate (Cliff & Ord 1973). As there was no significant spatial autocorrelation (Moran’s I statistic = −0.16, p = 0.69), spatial autocorrelation was not included in subsequent analyses.

The effects of landscape matrix and local grassland context on insect visitation frequency were analyzed using generalized linear mixed models (GLMMs). The response variable was insect visit frequency, and explanatory variables were forest edge length, landscape heterogeneity, field margin management, and those interaction terms (forest edge length × landscape heterogeneity, forest edge length × field margin management, and landscape heterogeneity × field margin management). The negative binomial distribution was used for the error term, and the field and observation dates were included as random terms. To detect effective variables and spatial scales, we performed model selection based on the Akaike information criterion (AIC) for all possible models with combinations of explanatory variables and spatial scales (171 models = 17 combinations of explanatory variables × 10 buffer radius sizes + null model). Effective variables in selected models were confirmed based on the absolute z-values (estimated coefficient / standard error) being greater than 2.0 (Burnham & Anderson 2002). Correlation coefficients between variables were never very large (|r| < 0.58), indicating no severe collinearity.

To examine the relationship between insect visitation frequency and the number of bacterial ASV_97_ and CFU, GLMMs and GLMs were conducted at flower and field scales, respectively. Response variables were the number of ASV_97_ per flower and field and CFU per flower, and explanatory variables were visitation frequency, the quadratic term of visitation frequency, and insect diversity calculated as the Simpson diversity index. Error terms were the Poisson and negative binomial distributions for the number of ASV_97_ and CFU, respectively. Fields and nectar sampling dates were used for random terms for the number of ASV_97_ per flower and CFU per flower. Additionally, because the number of ASV_97_ per flower and CFU has many zeros, zero-inflated components were included in GLMMs for these data. The zero-inflated component was only the intercept for the number of ASV_97_ per flower. For CFU per flower, because our pre-analysis had shown a significant relationship between visitation frequency and presence/absence of CFU, the zero-inflated component included visitation frequency, its quadratic term, and insect diversity. The statistical significance of each explanatory variable was tested using the likelihood-ratio test. Correlation coefficients between insect visit frequency and insect diversity were not large (r = −0.46), indicating no serious collinearity.

The differences in the number of bacterial ASV_97_ and in CFU among bagging treatments were analyzed using analysis of variance (ANOVA). Response variables, error terms, and random terms were the same as described above for the relationship with insect visit frequency, with bagging treatments as a fixed term. Zero-inflated components with only an intercept were also added to cope account for many zeros. Furthermore, to cope with heteroscedasticity among bagging treatments, variance parameters of each treatment were estimated in the same models. If results of ANOVA show a significant difference among bagging treatments, i.e., p < 0.05, multiple comparisons were performed using the Tukey-Kramer method as a post-hoc test.

To examine the relationship between nectar consumption volume and insect visitation frequency and to examine differences in nectar consumption volume among bagging treatments, linear mixed model (LMM) and ANOVA were performed, respectively. Nectar consumption volume was calculated as the difference in nectar volume that was collected from the same flower morphs (pin or thrum) and in the same fields and dates between the paper bag treatment and the other bag treatments (no, fine-mesh, and rough-mesh bag treatments). In LMM, nectar consumption volume was used as the response variable, and explanatory variables included insect visit frequency, insect diversity, and flower morphs (pin and thrum). Since thrum flowers tend to have more nectar than pin flowers (Cawoy et al. 2006), flower morphs were added as a covariate. The random terms were fields and sampling dates. Statistical significance of each variable was tested using the likelihood-ratio test. In ANOVA, nectar consumption volume was used as the response variable, and bagging treatments, excluding the paper bag treatment, and flower morphs were used as fixed variables. The random terms were the same as those described above for LMM. Additionally, to cope with heteroscedasticity among bagging treatments, variance parameters of each treatment were estimated in the same ANOVA model. If the results of ANOVA show a significant difference among bagging treatments, multiple comparison was conducted using the Tukey-Kramer methods as a post-hoc test. Furthermore, to determine whether nectar consumption volume was higher than zero, we conducted a t-test of the population mean.

GLMMs (LMM) and ANOVA were performed using glmmTMB package (Brooks et al. 2017), and likelihood-ratio test was conducted using car package (Fox & Weisberg 2019). AIC model selection was conducted using MuMIn package (Bartoń 2025). For multiple comparisons, multcomp package was used (Hothorn et al. 2008).

## Results

### Field observation

We observed a total of 1,960 insects visiting buckwheat flowers. The buffer radius size of the best and second-best models was 200 m according to AIC values, and ΔAIC (difference of AIC value from the best model) of the second model was 1.5 (**Figure 2A**). In the best model, insect visitation frequency was positively correlated with forest edge length (**Figure 2B, Figure S4, Table S1**). In addition, insect visitation frequency in fields with set-aside field margins was higher than that in fields with mown margins (**Table S1**). Nectar consumption volume was positively correlated with insect visitation frequency (**Figure 3A, Table S1**).

**Figure 3.**
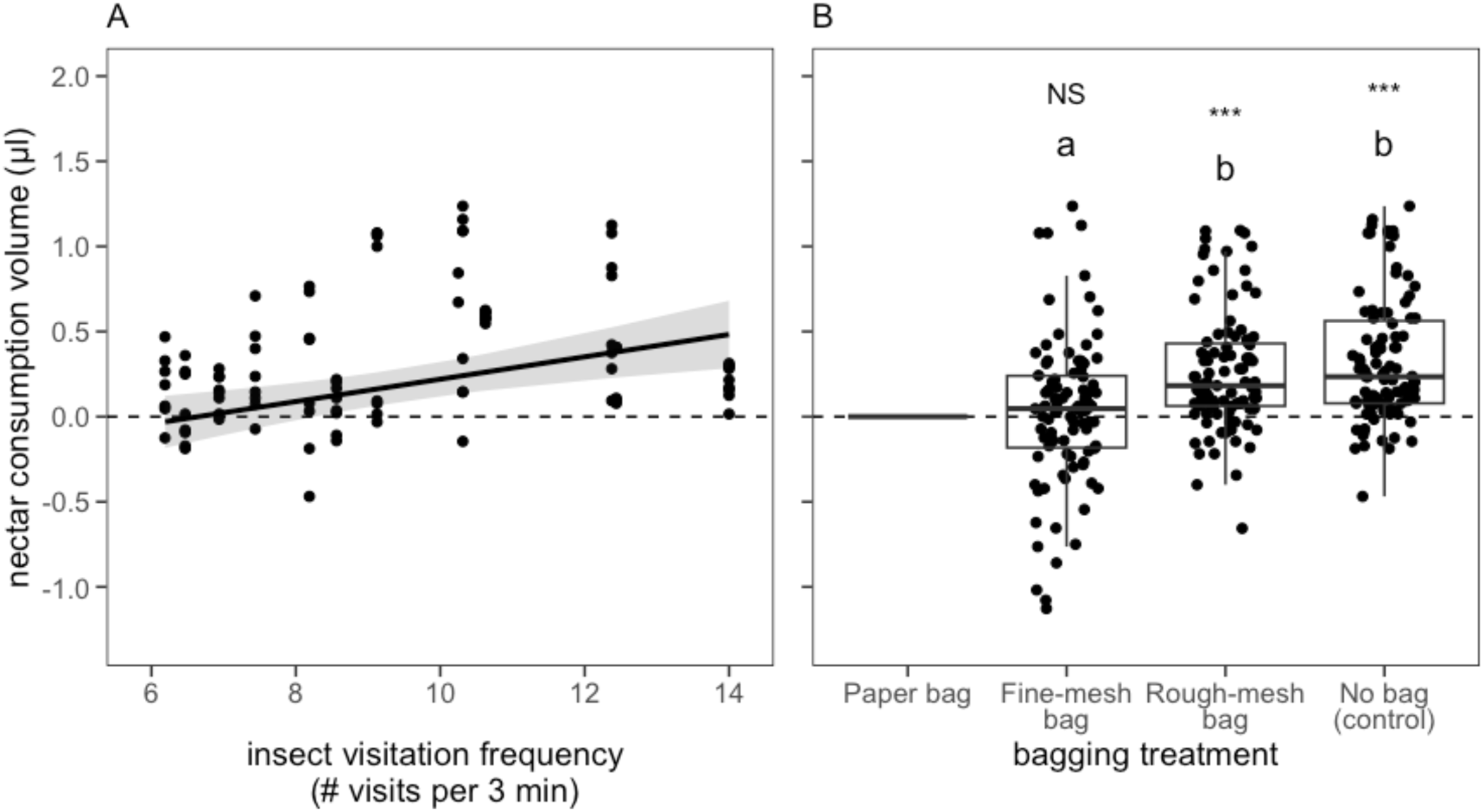
Results of field observation (A) and experiment (B) on nectar consumption by insects. (A) The estimated relationships between nectar consumption volume and insect visitation (mean number of insect visits observed per 3 minutes in a 1 x 10 m transect) (*R^2^* = 0.38, *p* < 0.001). (B) The effect of bagging treatment on nectar consumption volume. Different letters indicate significant difference (*p* < 0.05). Nectar consumption volume was calculated as the difference in nectar volume from flowers with the paper bag treatment. *NS* (*p* > 0.05) and *** (*p* < 0.001) indicate the results of a t-test that tested whether the mean of nectar consumption volume was larger than zero.

In the metabarcoding analysis, a total of 37 bacterial ASV_97_ belonging to 15 genera were identified (**Figure S3**). Taxa that are commonly found in the floral nectar environment (Quevedo-Caraballo et al. 2025), such as *Pseudomonas*, *Sphingomonas*, *Rosenbergiella*, and *Acinetobacter*, were frequently recorded in our samples (**Figure S3**). Both bacterial diversity (number of ASV_97_ per flower and per field) and abundance (log-transformed CFU per flower) showed hump-shaped responses to insect visitation frequency (**Figure 4A-C, Table S2**). The number of ASV_97_ and CFU per flower were negatively correlated, even when nectar volume was taken into consideration (partial correlation coefficients = −0.35, p = 0.004).

**Figure 4.**
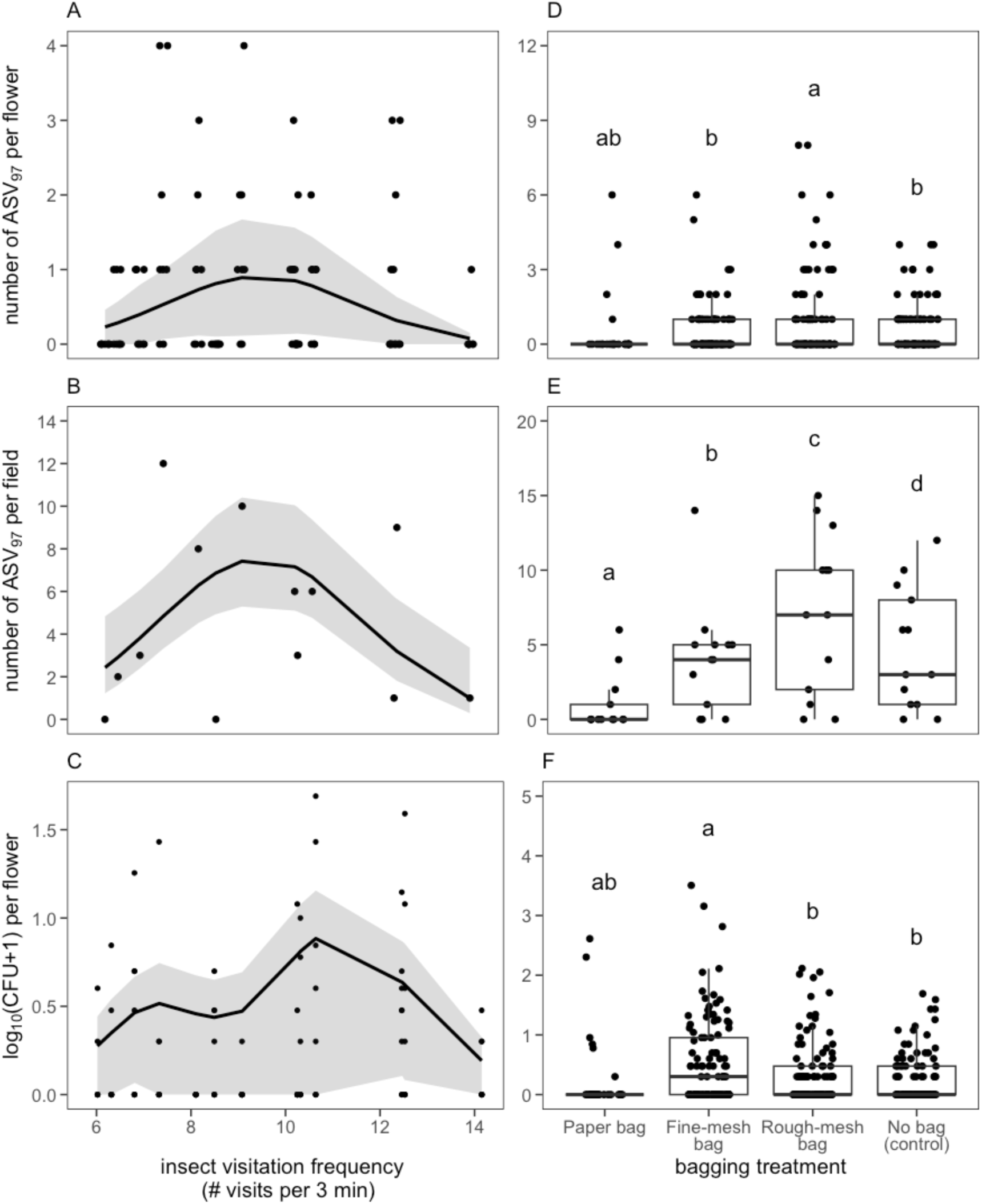
Results of field observation (A-C) and experiment (D-F). (A-C) The estimated relationships of bacterial diversity and abundance with insect visitation frequency. Estimated coefficients and statistical significances are shown in Table S2. (E-F) The effects of bagging treatment (Figure 1D) on bacterial diversity and abundance. Different letters indicate significant difference (*p* < 0.05).

### Field experiment

Consumption of nectar by insects was greater in the rough-mesh and no-bag treatments than in the fine-mesh bag treatment (**Figure 3B**, ANOVA, p < 0.001), as expected and consistent with the observational results (**Figure 3A**). Furthermore, nectar consumption volume in the rough-mesh and no-bag treatments was greater than zero, but not in the fine-mesh bag treatment (**Figure 3B**, t-test, fine-mesh, p = 0.96; rough-mesh, p < 0.001; no bag, p < 0.001), further confirming that insect visitation frequency varied among the bagging treatments.

In the metabarcoding analysis, 58 bacterial ASV_97_ belonging to 23 genera were identified (**Figure S3**), dominated by taxa typical of floral nectar such as *Pseudomonas*, *Sphingomonas*, *Rosenbergiella*, and *Acinetobacter* (Quevedo-Caraballo et al. 2025). Both bacterial diversity (number of ASV_97_ per flower and per field) and abundance (log-transformed CFU per flower) differed among bagging treatments (**Figure 4D-F**, ANOVA, ASV_97_ per flower, *p* = 0.007; ASV_97_ per field, *p* < 0.001; CFU per flower, *p* < 0.001). Diversity was higher in the rough-mesh treatment than in the fine-mesh and no-bag treatments (**Figure 3D, E**). Abundance in the fine-mesh bag treatment was higher than in the rough-mesh and no-bag treatments and did not differ from that in the paper-bag treatment (**Figure 3F**).

## Discussion

Taken together, our data indicate that landscape matrix indirectly determined the diversity of bacteria inhabiting buckwheat nectar via its effects on the frequency of insect flower visits (**Figure 2, 4**). Diversity peaked at the intermediate level of insect visitation frequency. This pattern was observed consistently at two spatial scales (flower and field) and in both observational and experimental data (**Figure 4**). In addition, as we were able to confirm that nectar consumption volume was positively correlated with insect visitation frequency, insect visitation frequency was likely a good proxy of bacterial dispersal rate (**Figure 3**). Therefore, our results support the intermediate dispersal hypothesis, which predicts the hump-shaped pattern of diversity that we found here with nectar bacteria. As such, this study links microbial diversity in buckwheat flowers, which are roughly 3 mm in radius, to the environmental conditions experienced by their vectoring animals over the comparatively vast scale of 200 m in radius within the rural landscape.

Theory suggests that the key mechanism driving the hump-shaped relationship between dispersal rate and species diversity is competitive exclusion (Cadotte 2006; Mouquet & Loreau 2003). According to this theory, the number of species in a local habitat patch generally increases as dispersal rate increases, but the probability that dominant competitors colonize in local habitat patches also increases. Consequently, competitive exclusion is predicted when dispersal is very frequent, resulting in the hump-shaped relationship. In our system, diversity and abundance per flower were negatively correlated, suggesting that competitive exclusion might have been responsible for the hump-shaped patterns. However, individual buckwheat flowers open for only a single day (Free 1993), which might be too short for competitive exclusion to play a large role. What might be an additional or alternative explanation for the observed hump-shaped patterns?

One possible explanation concerns the effect of insect visitation on bacterial extinction in addition to bacterial immigration. As insects introduce new bacteria to the flowers they visit, they also remove some of the bacteria already present in the nectar through nectar consumption. Thus, nectar-foraging insects contribute to both immigration and extinction of the microbes. However, the probability that new bacterial taxa are introduced into a flower probably increases rapidly at first and then begins to saturate as insect visitation frequency increases (Loreau & Mouquet 1999; Mouquet & Loreau 2003). Meanwhile, when a few insects visit the flower, the probability that all individuals of certain bacterial taxa present in the nectar are removed, i.e., a local extinction event, is expected to be low. However, a further increase in insect visitation frequency is likely to increase extinction rate exponentially, due to lower bacterial abundance. The combined result of this insect-driven immigration and extinction would be a hump-shaped relationship.

One way to conceptualize this hypothesis visually is to view bacterial diversity as a variable that is shaped by a dynamic balance between immigration and extinction, as illustrated in **Figure 5**. The equilibrium assumption made by this conceptual model (**Figure 5**) is unlikely to be met in real communities of nectar bacteria. However, knowing the equilibrium point to which these communities are attracted can still be informative in understanding diversity patterns, even when communities are in a non-equilibrium state (Fukami & Nakajima 2011; Roughgarden 1998). Of course, simple models like this should be treated with caution because they leave out other factors that may also affect diversity. For example, nectar consumption by insects may affect bacterial population size not just directly by removal from nectar, but also indirectly by altered temperature and oxygen availability in nectar.

**Figure 5.**
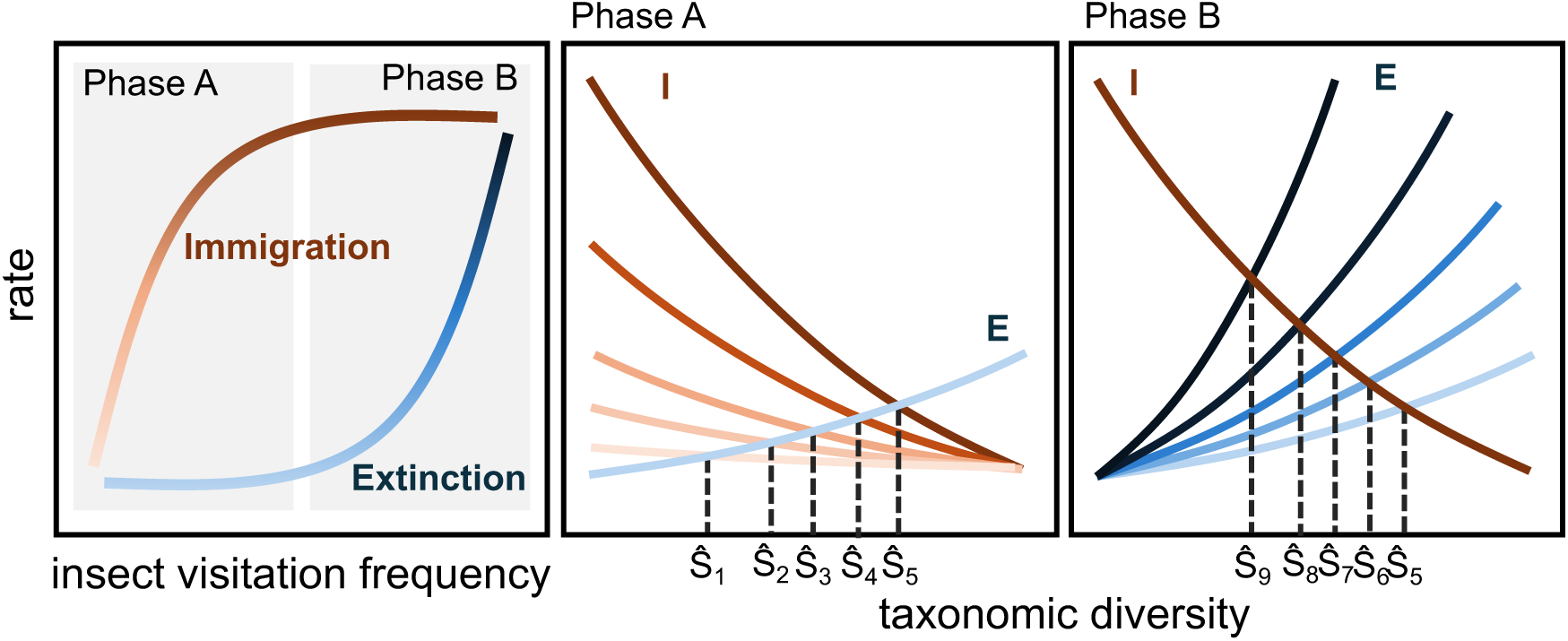
Hypothesized relationship between insect visitation frequency, the rate of immigration and extinction in nectar bacteria in flowers, and the taxonomic diversity of nectar bacteria in flowers. The symbol Ŝ_n_ denotes the predicted equilibrium level of taxonomic diversity, numbered from 1 to 9 in order of insect visitation frequency. This conceptual model predicts a hump-shaped relationship between insect visitation frequency and bacterial taxonomic diversity.

Hump-shaped patterns were observed in both diversity (**Figure 4A, B, D, E**) and abundance (**Figure 4C, F**). If competitive exclusion was the only reason for the diversity decline under high insect visitation frequency, we would not necessarily expect such a concomitant decline in abundance. On the other hand, we would expect hump-shaped patterns in both diversity and abundance if increased extinction accompanied increased immigration. It remains uncertain to whatr extent this dynamic balance between immigration and extinction explains the patterns we observed remains uncertain. However, similar phenomena seem possible whenever the habitat patches of the vectored organisms constitute consumable resources for the vectoring organisms. Besides nectar-inhabiting microbes dispersed by nectarivorous insects, other examples where this condition may apply include fruit-inhabiting microbes dispersed by frugivorous insects, carrion-inhabiting microbes dispersed by scavenging insects, and herbaceous plants dispersed by herbivorous mammals.

Given the results we present here, we suggest that future research can strengthen the rigor and scope of the inferences drawn in this study in three interrelated ways. First, it has become clear through this study that quantifying both immigration and extinction, not just immigration, can be important in understanding how vector-mediated dispersal affects diversity. We found no clear difference in bacterial taxonomic composition among the bagging treatments (**Figure S2, S3**), which suggests that no obvious differences may exist among bacterial taxa in their tendency to immigrate or go extinct and among insects in their vectoring bacterial taxa. However, it would be informative to more directly study which bacterial taxa insects tend to bring to nectar and which taxa they tend to remove from nectar. This work could involve examining bacteria in the mouthparts of flower-visiting insects (Warren et al. 2025) before and after flower visits.

Second, this study quantified insect visitation frequency rather coarsely by looking at overall frequency without considering the taxonomic identity of the insects. We did make use of the diversity of buckwheat-visiting insects by applying bags of different mesh sizes to manipulate visitation frequency according to their size. However, studying the contribution of different insects to microbial dispersal in more detail could reveal more nuanced dispersal–diversity relationships than were possible in this study. Experimental exclusion of specific insect groups would help for this purpose. Exclusion is feasible with ants, for example, which are major pollinators of buckwheat (Natsume et al. 2022).

Finally, this study did not consider the variation among insect species in their foraging distance, which can be substantial, with flightless arthropods such as ants and mites moving over just a few meters or less over their entire lives (Rhainds & Shipp 2004), while bees, beetles, butterflies, and other winged insects move across several hundred meters or more on a daily basis (e.g., Danforth et al. 2019). These insects are likely to contribute to microbial dispersal among flowers at greatly different spatial scales. Using the terminology of Fukami (2005), short-distance movement contributes to internal dispersal (i.e., dispersal of species across local communities within a metacommunity, a la Leibold et al. 2004). In our case, this scale may correspond to dispersal within a buckwheat field. In contrast, long-distance movement contributes to external dispersal (i.e., supply of species to local communities from the regional species pool, unaffected by the population dynamics happening within metacommunities, a la MacArthur & Wilson 1967). Theoretical work suggests that internal and external dispersal can interactively, rather than additively, influence diversity (Fukami 2005). Future studies that examine how different insect groups contribute to microbial dispersal at various spatial scales could provide a deeper understanding of vector-mediated dispersal. Although limited in scope to draw firm conclusions at this stage, a preliminary analysis of the data we present in this paper suggests that small- and large-bodied insects, which likely contribute more to internal and external dispersal, respectively, determine diversity in an interactive, rather than additive, manner (**Figure S5**).

In conclusion, our results have shown that landscape context can shape diversity patterns in vector-dispersed organisms by affecting their vectors. We have discussed the possibility that the intermediate dispersal pattern observed may readily emerge in vector-dispersed organisms even without competitive exclusion playing a large role, if the vectors affect both immigration and extinction. Although less studied than vector-independent dispersal, vector-mediated dispersal is a common phenomenon not just in bacteria, but also in plants, animals, and fungi (e.g., Borgmann-Winter et al. 2023; Lovas-Kiss et al. 2020; Plue & Cousins 2018; Soons et al. 2016; Suetsugu et al. 2018). Given that these two types of dispersal may be governed by different constraints across landscapes, we suggest that more research on vector-mediated dispersal would lead to a better understanding of how dispersal affects biodiversity.

## Acknowledgments

We are grateful to Tadashi Miyashita for guidance and support for our field surveys and comments on the manuscript; Hisao Saito for support for field surveys; and Hongo Nousan Ltd., Tagiri Nousan Ltd., and all landowners for permitting us to do this work in their fields. We thank to Hisatomo Taki for lending us his rough-mesh bags. We also thank Hidenori Deto and Yen-Hua Yeh for support for field surveys and Jessica Aguilar, Rosa McGuire, Andrea Nebhut, and other members of the Community Ecology Group at Stanford University for comments. This study was funded by JSPS KAKENHI (grant number: 23K27246, 24K01777), JST FOREST (Fusion Oriented REsearch for disruptive Science and Technology, grant number: JPMJFR2247), and the Stanford Doerr School of Sustainability Discovery Grants.

## Notes

### Competing Interest Statement

The authors have declared no competing interest.

