## Supplemental Text for "Diversity of vector-dispersed microbes peaks at a landscape-defined intermediate rate of dispersal"

**Supplemental Text 1**

On September 5-7, 2019, we conducted a preliminary study of the prevalence and abundance of bacteria and yeasts in buckwheat nectar. We collected buckwheat nectar from 12 fields in the same area as for the main study in 2020. In each field, we sampled nectar from eight plants (four pin and four thrum), except in two of the 12 fields where we sampled nectar from either 10 or 16 plants, for a total of 106 samples. At each field, the plants were chosen haphazardly along the field margin. From each of these plants, we extracted nectar from four flowers belonging to the same inflorescence using a 0.5-µl microcapillary tube, between 11:00am and 5:30pm. We placed the nectar collected from the four flowers into a separate PCR tube containing 40 µl of distilled water. The PCR tubes with diluted nectar samples were kept cool in a Styrofoam box with ice packs in it until transported to the laboratory. Within two hours of the sampling in the field, 10 µl of each diluted sample were plated onto a TSA plate with cycloheximide to obtain CFU (colony forming units) counts. For each sample, 10 µl of the remaining sample was plated onto a YM plate with chloramphenicol. We did the CFU counting two and seven days after plating for TSA and YM, respectively. None of the 106 nectar samples yielded more than 9 yeast-like CFU, with 61 (58 %) of the plates yielding none, on the YM plates. Many more CFU were observed on the TSA plates, with 3 (3%), 18 (17%), 48 (45%), and 37 (35%) plates yielding >100, 10-100, 1-10, and no CFU, respectively. This preliminary result indicates that bacteria are frequently found, often in high abundance, whereas yeasts are rare, in buckwheat nectar in our study landscape.

**Supplemental Text2**

DNA metabarcoding

Using the nectar samples obtained, we also conducted microbial DNA extraction, PCR, and amplicon sequencing to examine bacterial taxonomic composition. We extracted DNA using the NucleoSpin 96 Tissue (Macherey-Nagel, Dueren, Germany). The extraction followed the protocol specified by the manufacturer, but with some modifications. Specifically, 20 µl of each thawed sample (and PCR-grade water as negative control) was combined with 4 µl of zymolyase enzyme (MP Biomedicals, Burlingame, CA, USA) within a microcentrifuge tube. These mixtures were incubated at 30°C for 30 minutes. Once the incubation period was complete, we lysed the samples. The samples were then incubated at 56°C overnight (14 hours). The next morning, we did DNA binding. We followed the protocol as stated in the Genomic DNA from Tissue User manual for NucleoSpin® 96 Tissue 2014, with the only exception being that the final elution of DNA was done with 60 µl of Buffer BE (Macherey-Nagel) rather than 100 µl to increase the final DNA concentration in the eluate. The final extracted DNA from each sample was stored in a sterile microcentrifuge tube at 4°C until library preparation. Bacterial amplicon sequencing involved the same methods as in Warren et al. (2025), using primers 515F (Parada) and 806R (Apprill) to amplify the highly variable (V4) region of the 16S rRNA gene (Apprill et al. 2015; Caporaso et al. 2012; Parada et al. 2016; Warren et al. 2025).

Bioinformatics

All bioinformatics steps were conducted using R. The Divisive Amplicon Denoising Algorithm 2 (DADA2, Callahan et al. 2016) pipeline was used to merge paired-end sequences, quality filter and trim these sequences, remove chimeras, and cluster sequences into amplicon sequence variants (ASV). Taxonomy of each ASV was assigned using the SILVA version 138.1 database (Yilmaz et al. 2014). To minimize grouping ASV from the same genome into separate clusters, ASV were re-clustered into sequences with a 97% similarity threshold using the DECIPHER (Wright 2016) and speedyseq packages (McLaren 2025). We will refer to these re-clustered sequences as ASV_97_. Those ASV_97_ that were detected from DNA extraction, PCR, water negative controls were removed from nectar samples before data analysis. We assumed that the probability of cross-contamination from nectar samples to negative controls was higher in the outdoor field setting than in the laboratory. For this reason, we did not remove any ASV_97_ observed in field negative controls from nectar samples. Cyanobacteria, Rickettsiales, and *Cellulosimicrobium* were removed because these taxa were clearly not expected to be found in nectar. Singleton ASV_97_ that were present in only a single sample were removed from nectar samples. Finally, to confirm the sufficiency of our DNA metabarcoding results, rarefaction analysis was conducted. The observed numbers of ASV_97_ for all nectar samples were in the range of saturation with respect to the read number (**Figure S1**).
