## Supplemental Figures & Tables for "Diversity of vector-dispersed microbes peaks at a landscape-defined intermediate rate of dispersal"

**Supplementary Figures and Tables**

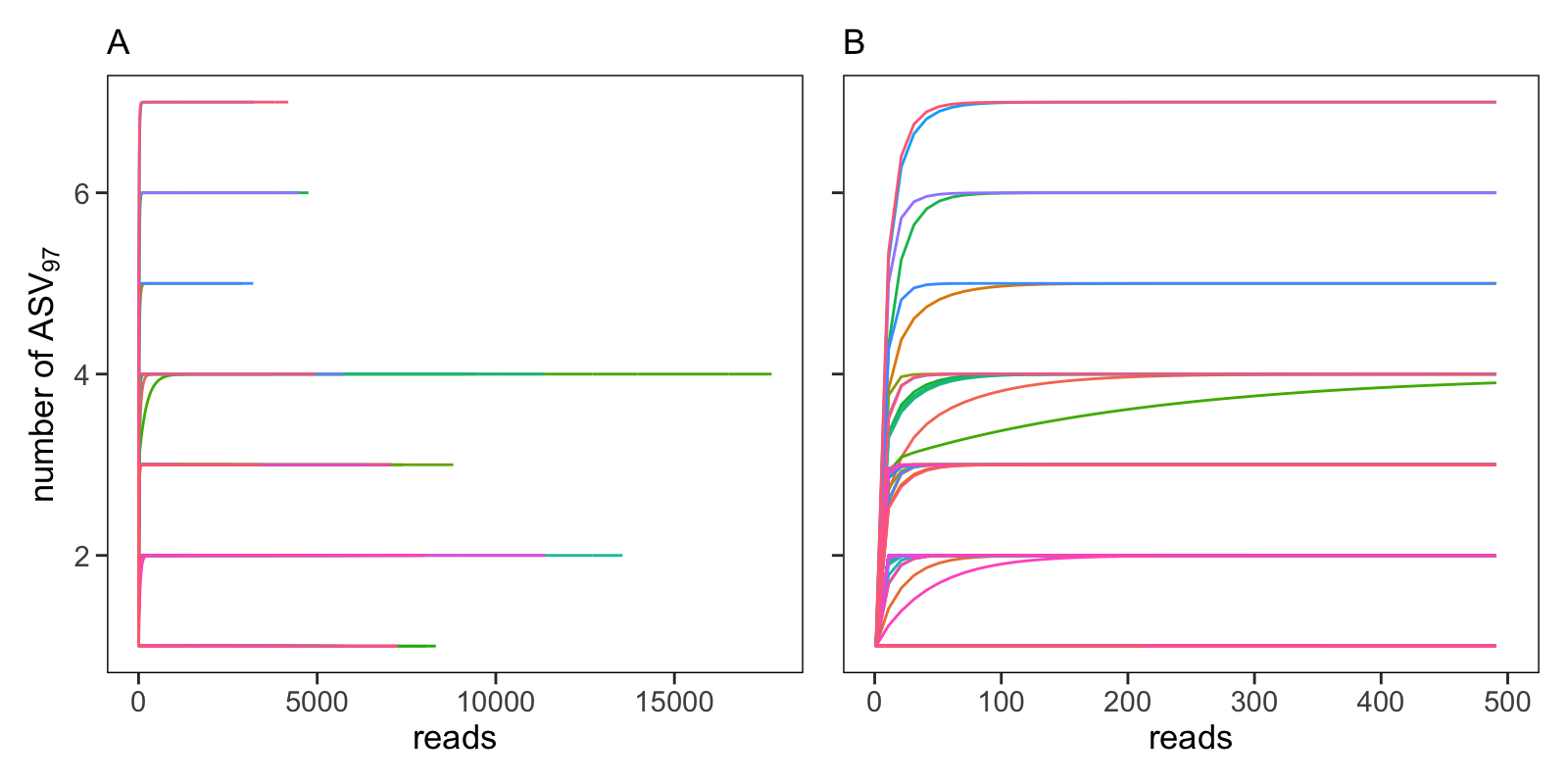

**Figure S1**	The results of rarefaction analysis. (A) results for whole reads and (B) results for up to 500 reads. Rarefaction analysis was conducted using vegan package in R (Oksanen et al. 2025).

Oksanen, J., G. L. Simpson, F. G. Blanchet, R. Kindt, P. Legendre, P. R. Minchin, et al. 2025 vegan: community ecology package. R package version 2.8-0.

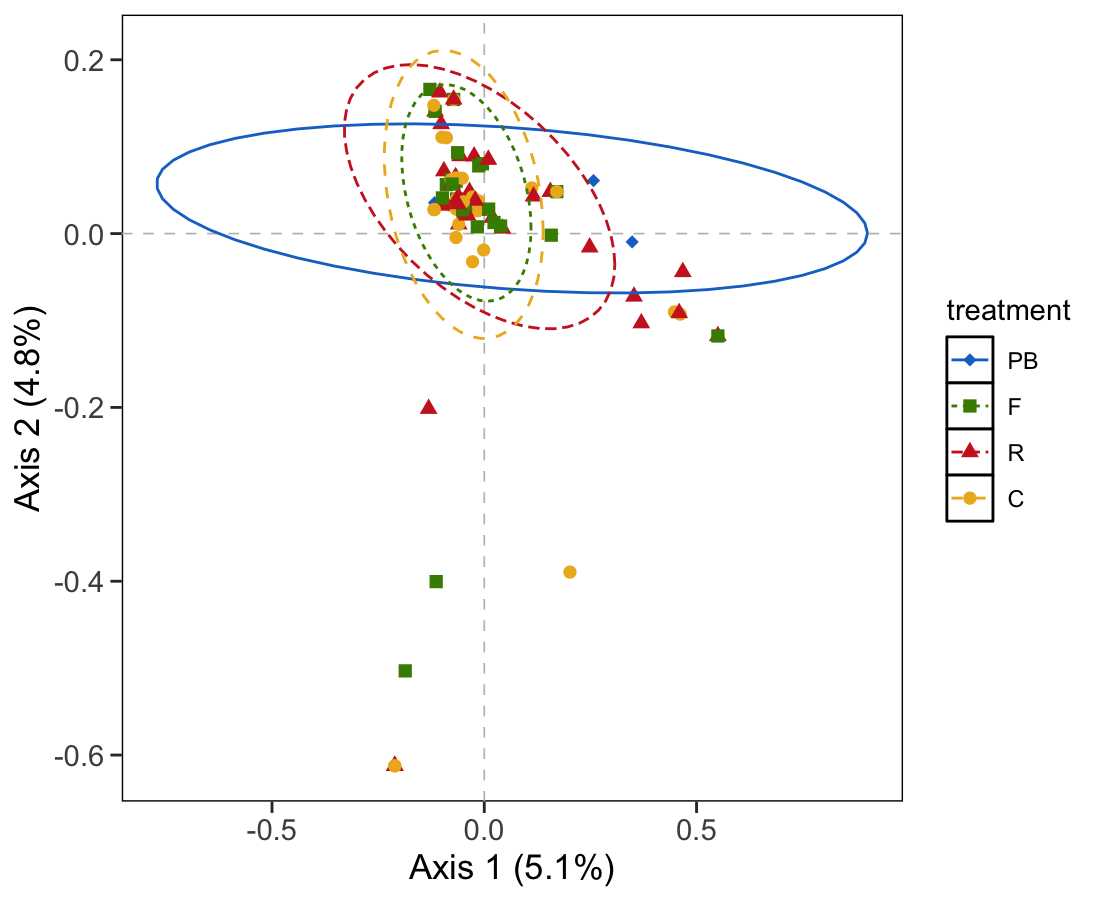

**Figure S2** Biplot of bacterial communities relative to the first and second PCoA (principal coordinate analysis) axes using the Jaccard dissimilarity index in each of the bagging treatments. The points indicate the bacterial communities of each buckwheat flower. PB: paper bag, F: fine-mesh bag, R: rough-mesh bag, and C: control (no bag). PCoA and PERMANOVA were conducted using ape (Paradis & Schliep 2019) and vegan (Oksanen et al. 2025) packages, respectively.

Oksanen, J., G. L. Simpson, F. G. Blanchet, R. Kindt, P. Legendre, P. R. Minchin, et al. 2025 vegan: community ecology package. R package version 2.8-0.

Paradis, E., and K. Schliep. 2019. ape 5.0: and environment for modern phylogenetics and evolutionary analyses in R. Bioinformatics, 35: 526–528.

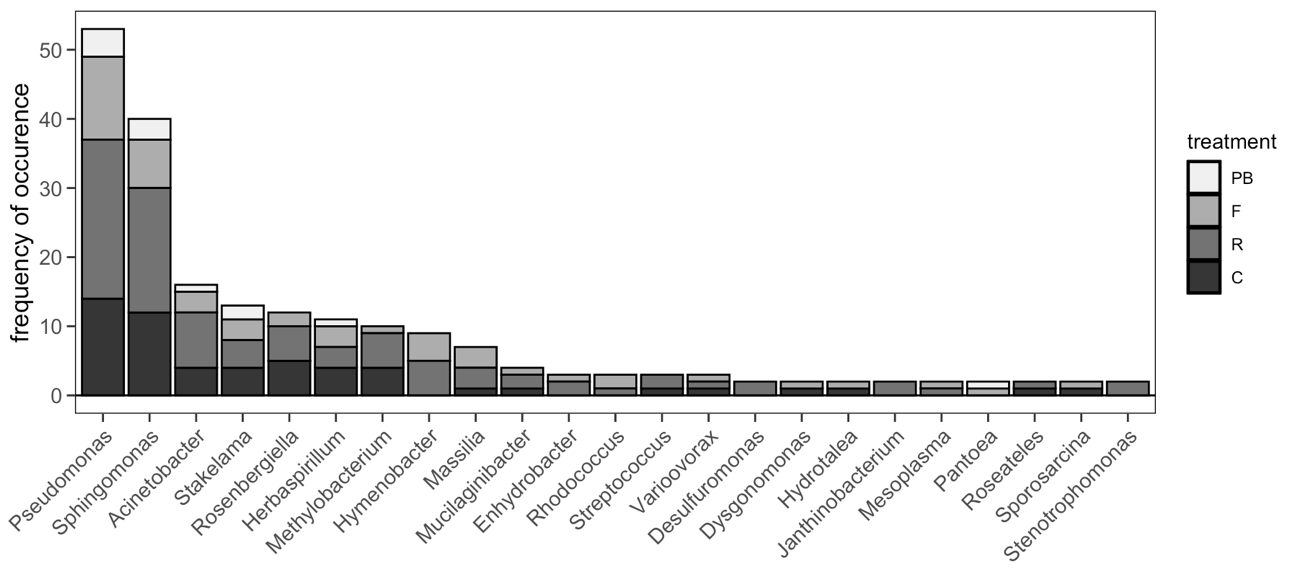

**Figure S3**	The frequency of occurrence of each bacterial genus. PB: paper bag, F: fine-mesh bag, R: rough-mesh bag, and C: control (no bag)

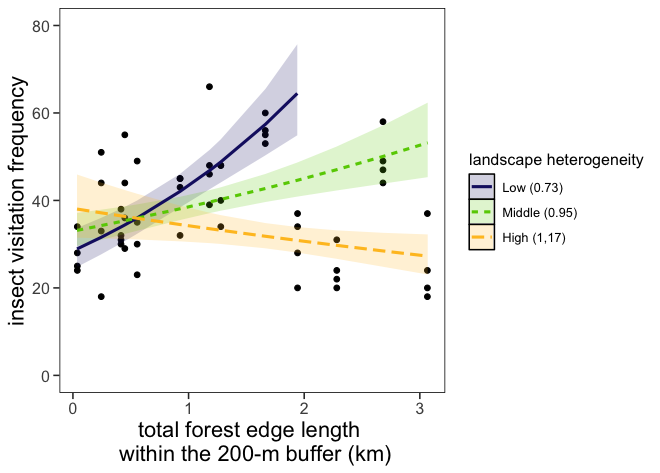

**Figure S4**	Relationship between landscape characteristics (as quantified by forest edge length and landscape heterogeneity) and the frequency of insect visits to buckwheat flowers. Data points represent four observations at each of the 13 study fields. Because landscape heterogeneity was positively correlated with the proportion of buckwheat fields (r = 0.51), the negative relationship between landscape heterogeneity and insect visit frequency was likely caused by dilution of insect among surrounding buckwheat fields. Insects might be diluted and less abundant in buckwheat fields with many surrounding buckwheat fields, even though there is a longer forest edge where is potentially one of the main habitats of insects.

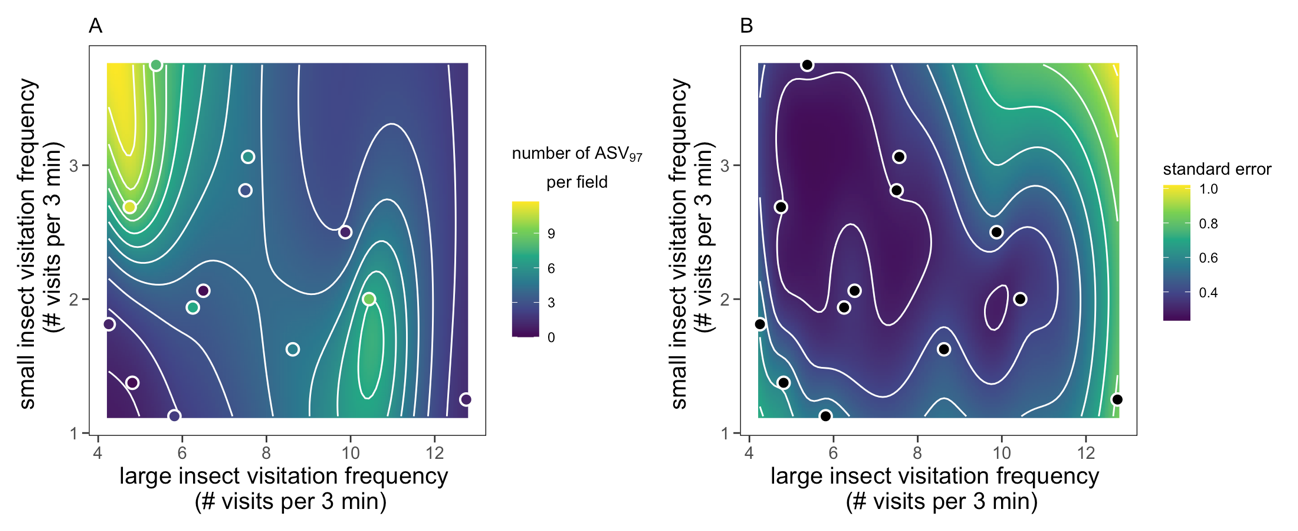

**Figure S5**	The estimated relationship between the number of bacterial ASV_97_ per field and the frequency of small and large insect visits; (A) estimated value and (B) estimated standard error. Each dot represents each field. This relationship was estimated using a generalized additive model (GAM) with the REML method (restricted maximum likelihood estimation) to avoid overfitting. The response variable was the number of ASV_97_ per field, and explanatory variables (smooth terms) were large and small insect visit frequency and the interaction term. The Poisson distribution was used for the error term. The smooth component of the interaction term was statistically significant (GAM, *r^2^* = 0.3, proportion of variance explained = 63.1%, Wald test, p = 0.009). This analysis was conducted using the mgcv package (Wood 2017).

Wood, S. N. 2017. *Generalized additive models: an introduction with R*. Second edition. Chapman and Hall.

**Table S1**	Results of AIC model selection. The values in each column indicate z-value (estimate / S.E.). FEL: forest edge length, LH: landscape heterogeneity, and FMH: field margin management. This table shows the results of top five models.

| **AIC** | **∆AIC** | **Radius size** | **FEL** | **LH** | **FMM** | **FEL × LH** | **FEL × FMM** | **LH × FMM** |
| --- | --- | --- | --- | --- | --- | --- | --- | --- |
| 377.6 | 0 | 200 | 4.00 | -6.44 | 3.66 | -5.97 | - | 3.05 |
| 379.1 | 1.5 | 200 | 3.56 | -6.48 | 3.69 | -5.66 | -0.70 | 3.13 |
| 382.5 | 4.9 | 250 | 4.92 | -6.60 | 2.25 | -5.73 | -2.67 | 4.44 |
| 384.6 | 6.5 | 200 | 3.07 | -5.09 | 3.20 | -4.39 | - | - |
| 386.6 | 8.4 | 200 | 2.35 | -5.10 | 3.20 | -4.08 | 0.28 | - |

**Table S2**	Results of GLMMs (GLMs and LMM) and likelihood-ratio test for the relationship between insect visit and nectar consumption volume, the number of bacteria ASV_97_, and CFU.

| **Response variables** | ***R^2^*** | **Explanatory variables** | **Estimate ± S.E.** | ***p*** |
| --- | --- | --- | --- | --- |
| Nectar consumption volume | 0.38 | Insect visitation frequency | 0.07 ± 0.02 | < 0.001 |
|  |  | Insect diversity | 3.08 ± 1.06 | 0.004 |
|  |  | Flower morph | 0.31 ± 0.06 | < 0.001 |
| Number of ASV_97_ per flower | 0.19 | Insect visitation frequency | 2.13 ± 1.07 | 0.05 |
|  |  | Insect visitation frequency^2^ | -0.11 ± 0.06 | 0.05 |
|  |  | Insect diversity | -5.90 ± 7.03 | 0.40 |
| Number of ASV_97_ per field | 0.55 | Insect visitation frequency | 1.86 ± 0.68 | 0.001 |
|  |  | Insect visitation frequency^2^ | -0.10 ± 0.03 | 0.001 |
|  |  | Insect diversity | -4.88 ± 3.47 | 0.16 |
| CFU per flower | 0.22 | Insect visitation frequency | -1.98 ± 1.50 | 0.33 |
|  |  | Insect visitation frequency^2^ | -4.61 ± 2.74 | 0.05 |
|  |  | Insect diversity | -0.51 ± 0.58 | 0.84 |
